# Fast and accurate bootstrap confidence limits on genome-scale phylogenies using little bootstraps

**DOI:** 10.1101/2021.07.21.453255

**Authors:** Sudip Sharma, Sudhir Kumar

**Author notes:** **Corresponding author: Sudhir Kumar**, Temple University.

## Abstract

Felsenstein’s bootstrap resampling approach, applied in thousands of research articles, imposes a high computational burden for very long sequence alignments. We show that the bootstrapping of a collection of little subsamples, coupled with median bagging of subsample confidence limits, produces accurate bootstrap confidence for phylogenetic relationships in a fraction of time and memory. The little bootstraps approach will enhance rigor, efficiency, and parallelization of big data phylogenomic analyses.

---

The bootstrap approach, introduced more than 35 years ago by Joseph Felsenstein^1^, has been the standard method to place confidence limits on inferred molecular phylogenies^2^ (Fig. 1a). If a group of sequences is recovered in a large proportion of bootstrap phylogenies (bootstrap confidence limit, BCL), their evolutionary relationship is considered well‐supported^1,3^. The bootstrap (BS) approach is being applied to increasingly larger datasets due to the widespread accessibility of genome sequence databases and the assembly of multispecies and multigene alignments containing hundreds of thousands of bases (e.g.,^4–6^). These large datasets have the power to reconstruct hard‐to‐resolve evolutionary relationships with high confidence (BCL > 95%)^4,5,7–9^, but they impose increasingly onerous computational demands because the computational complexity of phylogenomic analyses using the maximum likelihood (ML) method increases exponentially with the number of sequences and linearly with sequence length^10^. Consequently, the standard BS resampling procedure can take days to complete for big datasets^5,10^. Many heuristics have been proposed to moderate the escalation due to the increasing number of sequences (e.g., ref.^10,11^). However, no effective approaches are available to deal with the onerous computational burden imposed by an increase in sequence length due to the widespread adoption of next‐generation sequencing methods. Thus, the standard BS approach’s computational burden has become a new bottleneck in ensuring robust and reproducible phylogenomic analyses^12,13^.

**Figure 1.**
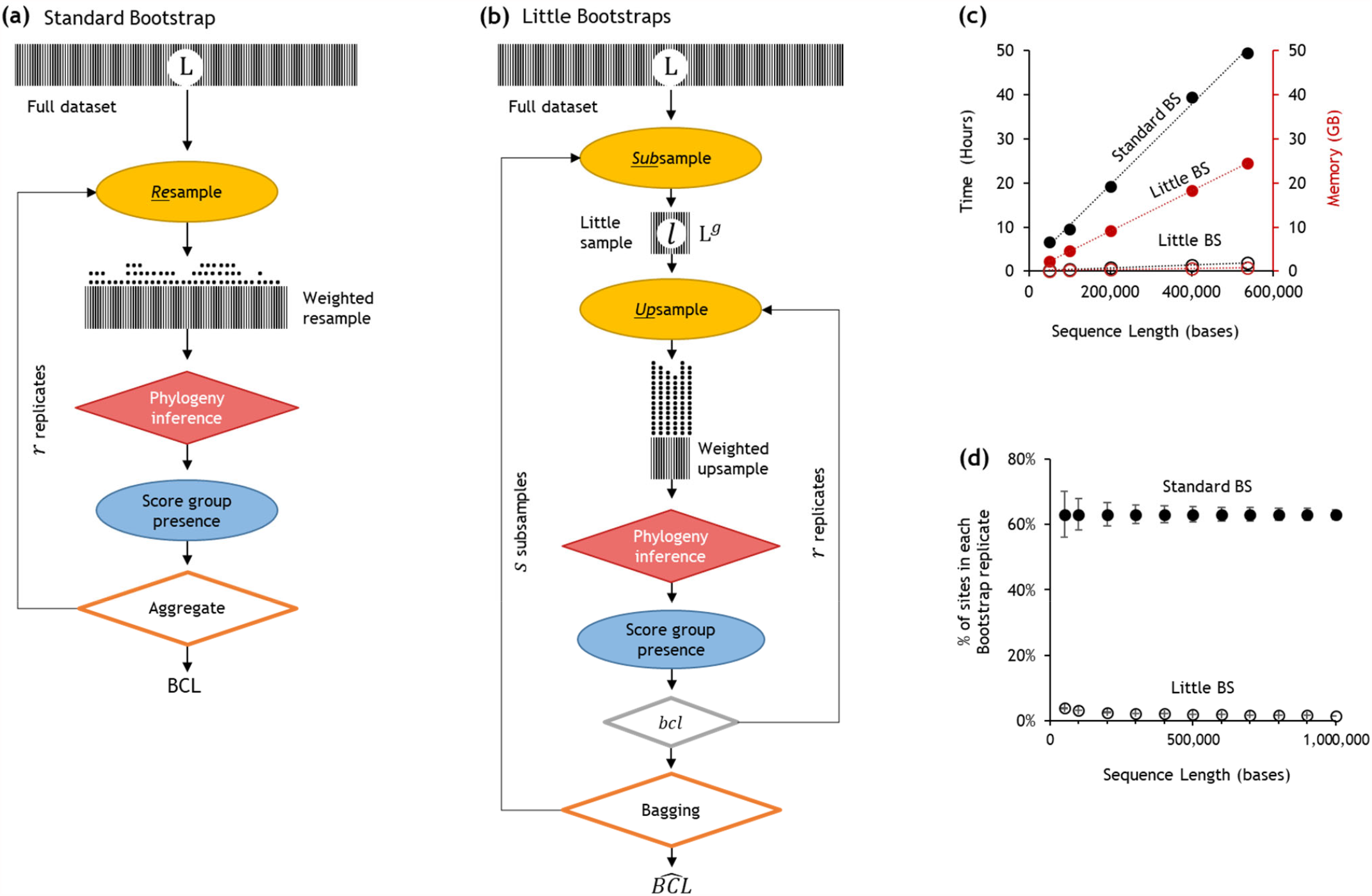
Overview of the Little Bootstraps approach. | Steps of **(a)** the standard phylogeny bootstrap^1^, and **(b)** the bag of little bootstraps (BS) approach. Shaded boxes represent sequence alignments in which denser hatching corresponding to a larger number of site configurations. The width of the box represents the sequence length. The generation of bootstrap replicate datasets differs between standard and little BS. In standard BS, L sites are randomly sampled with replacement from the original dataset containing L sites. In this *resampling* process, ∼63.2% of the data points^14,20^ are expected to be represented in a bootstrap replicate dataset^14^. Each replicate dataset is compressed into weighted resamples that contain only distinct site configurations and a vector of their counts (represented by stacks of dots). The ML tree is inferred from each replicate dataset and the ***BCL*** for a species group is the proportion of times that appeared in bootstrap replicate phylogenies. In little BS, L sites are randomly sampled with replacement from the little dataset consisting of only ***l*** = L^*g*^ sites to build each replicate dataset. Because ***l*** ≪ L, each site will be represented many times in the little bootstrap replicate dataset, which we refer to as *upsampling*. Upsampling only alters the frequency of unique site configurations. Stacks of dots are much higher for little BS due to upsampling than for standard BS that involves only resampling. The number of distinct site configurations in the upsampled dataset is smaller than that in the standard bootstrap replicate dataset, because ***l*** ≪ L. Therefore, ML phylogeny for little BS replicates is expected to require less time and memory, as long as is less than 0.632L on average. **(c)** Time and memory savings per replicate of little bootstrap (open circles) compared to the standard bootstrap (closed circles) for large datasets. Simulated dataset contained 446 taxa and sequence length ranges from 50,000 to 536,534. **(d)** The proportion of sites included in the bootstrap replicates for little datasets with ***l*** = L^0.7^ (open circles) and standard bootstrap (closed circles). The choice of ***l*** = L^0.7^ offers increasingly greater computational savings for longer sequences because of a decreasing proportion of sites included in the little samples. For example, the little dataset size is ∼3.1% of the original alignment for L = 100,000 bases, but it decreases to ∼1.6% when L increases 10‐fold (1,000,000 bases). Overall, memory and time savings greater than ∼95% can be achieved for phylogenomic data with long sequences.

## RESULTS AND DISCUSSION

### The little bootstraps (BS) approach for phylogenomics

Kleiner et al. proposed a bag of little bootstraps^14^ (little BS) approach to overcome statistical limitations of divide‐and‐conquer approaches^14–16^. Here, we introduce this BS approach for placing confidence limits on molecular phylogenies inferred using sequence alignments. In the little BS approach, bootstrapping is performed independently on ***s*** little datasets, each containing *l* sites sampled randomly without replacement from the full dataset with L sites (*l* ≪ L). A bootstrap confidence limit (*bcl*_*i*_) is estimated for each little dataset *i* by generating *r* phylogenies from bootstrap resampled datasets (Fig. 1b).

In little BS, the bootstrap resampling of little sample alignments is different from that of the standard BS, as L sites are sampled with replacement from *l* sites of the little subsample to build replicate datasets. Because *l* ≪ L, the same site is selected many times (up‐sampling) to build the bootstrap replicate dataset (Fig. 1b). A replicate phylogeny is estimated for each little BS replicate dataset. Then, the bootstrap confidence limit 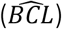 for a given group of species is derived from ***s*** little sample *bcl* values^14^, a procedure referred to as bagging. The average of ***s*** little sample *bcl* values, called mean‐bagging 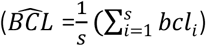 was found to work well in general statistical analyses, including computer‐simulated datasets^14^.

In the little BS approach, every site of the little sample is included, on average, L/ *l* times in the bootstrap replicate dataset. As these replicate datasets have the same number of sites as the full dataset, it obviates *ad hoc* corrections needed in other divide‐and‐conquer approaches and has desirable asymptotic theoretical properties^14–16^. The computational burden of ML phylogeny estimation is proportional to the number of distinct site configurations, so each little BS replicate’s time and memory requirements are of order *O* (L/ *l*) needed for a standard BS replicate. Kleiner et al.^14^ have suggested that little samples of size *l* = L^*g*^ (0.5 < *g* < 1.0) can reduce time and memory by orders of magnitude. In phylogenomics, these savings can be substantial (Fig. 1c) and grows as the length of the sequence alignment increases from thousands to millions of sites for a given value of the power parameter ***g*** (Fig. 1d).

### Performance of little BS for a computer‐simulated dataset

Simulations are frequently used to test the accuracy of computational phylogenetic methods because the true evolutionary relationships are known^17,18^ and used as the ground truth. So, we first present results of ML phylogenetic analysis of a computer‐simulated alignment containing 446 species and 134,131 sites (Fig. 2a). We conducted 100 standard BS replicates, an *ad hoc* convention adopted in many studies to make calculations feasible (e.g., ref.^12^). It required 6.1 GB of memory and 13.1 CPU hours per replicate (54 CPU days of total computation). These analyses established the true evolutionary relationships among sequences with very high confidence, i.e., *BCL* ≥ 95% for all 443 correct species groupings.

**Figure 2.**
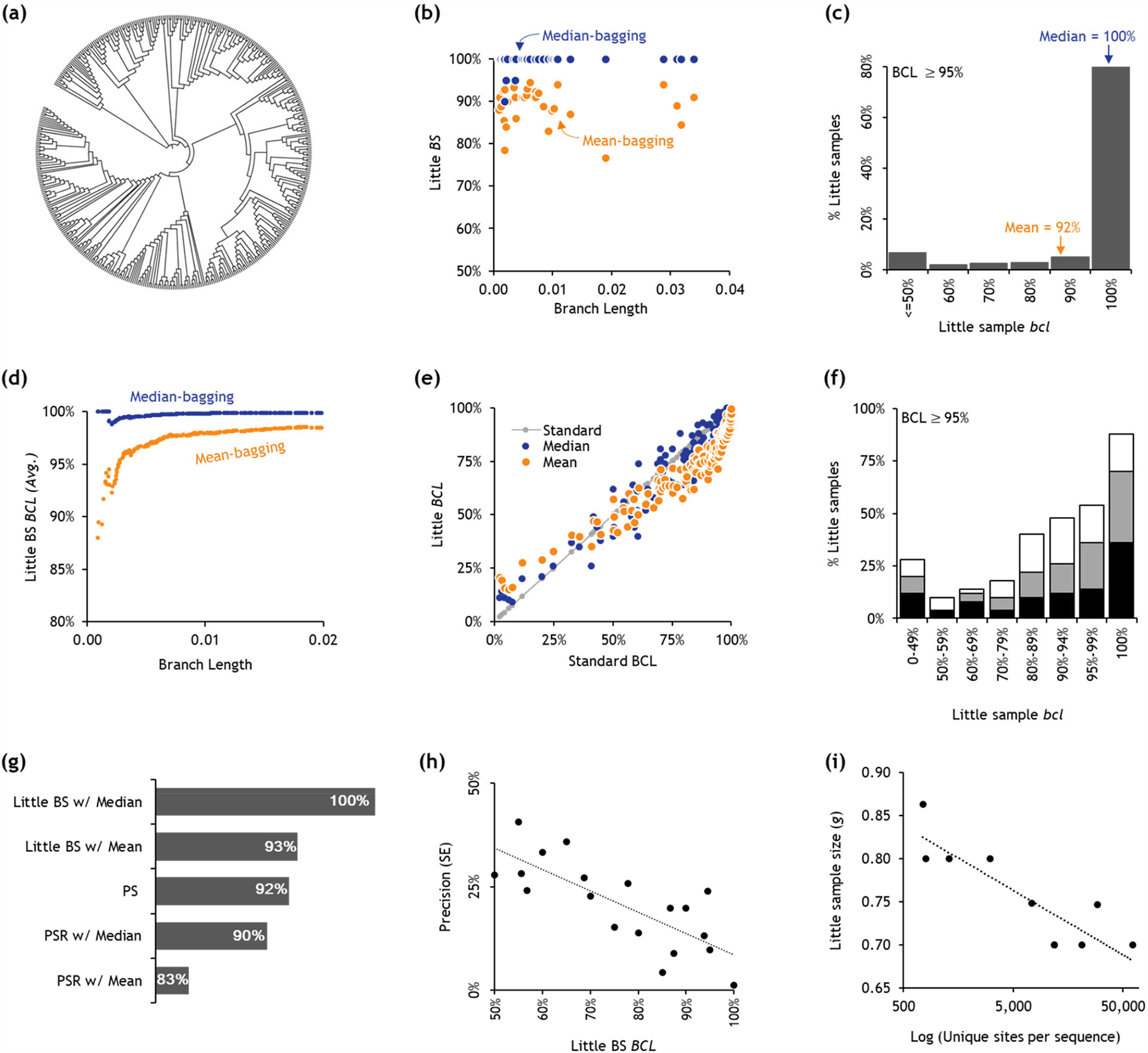
Little BS analyses of simulated and empirical phylogenomic datasets. | **(a)** A model phylogeny of 446 species based on the bony‐vertebrate clade from the Timetree of Life (See *Methods* section), which was used for simulating sequence evolution. A sequence alignment for 100 genes was generated in which evolutionary rates varied extensively among genes and evolutionary lineages following biologically realistic parameters and models (see *Methods* section). **(b)** The relationship of branch lengths and 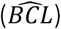 produced by little BS with mean‐bagging (orange) and median‐bagging (blue) for *l* = L^0.7^. The x‐axis is restricted up to the branch length of 0.04 because 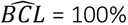 for mean and median bagging for longer branches. **(c)** The distribution of *bcl*_i_s for 49 species groups that received 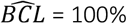 in little BS with mean‐bagging analysis of large datasets. **(d)** The average 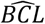 for all the species groups connected to the phylogeny with a given cutoff branch length (x‐axis). The x‐axis is restricted to 0.02 because mean‐ and median‐bagging performance does not change any further for longer branches. **(e)** The relationship of standard BS (*BCL*) and little BS 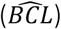 with mean‐bagging (orange circles) and median‐bagging (blue circles) for datasets smaller than 10,000 sites (***l*** = L^0.9^). The gray line shows the 1:1 relationship with the standard BS. The little BS offered time savings up to 37% and memory savings up to 42% in these small data analyses. The linear regression slope is 0.97 (*R*^2^ = 0.93) for median‐bagging and 0.89 (*R*^2^ = 0.89) for the mean‐bagging. However, a second‐order polynomial fits the mean‐bagging results better (*R*^2^ = 0.93). **(f)** The distribution of little sample *bcl*s for species groups in smaller datasets for which standard BS ***BCL*** ≥ 95% (black bars = 9,359 sites, gray bars = 7,002 sites, and white bars = 4,070 sites). **(g)** The true positive rates (TPR) for little BS with mean‐ and median‐bagging compared to other phylogenomic subsampling approaches (PS and PSR with Mean and with Median) in which upsampling was not applied (see *Methods* section). **(h)** The relationship of subsample size (*g*) and the number of unique site configurations per sequence (*C*) in empirical datasets. The log‐linear regression slope is ‐0.032 (*R*^2^ = 0.76). **(i)** The relationship between the little BS 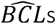 and their precision (standard errors, SEs) for the selected little BS parameters (Table 1). The linear regression slope is ‐0.52 (*R*^2^ = 0.59).

For the 446×134,131 dataset, we generated 10 little samples (*s* = 10) containing *l* = L^0.7^ sites (3,884 sites), with ten little BS replicates each (*r* = 10). ML phylogeny inference of these 100 little datasets required, on average, only 0.3 GB RAM and 0.6 hours of CPU time, offering a 95% reduction in memory and in time compared to the standard BS. With these improvements in efficiency, many little BS datasets could be run concurrently on a multicore desktop with 8 GB of RAM, unlike the standard bootstrap analyses that took up almost all the memory for estimating the ML phylogeny for one replicate dataset.

**Table 1.** Little bootstrap analysis of empirical datasets of varying length

| Species<sup>a</sup> | Full Dataset |  |  | Little BS subsamples |  |  |  |  |  | Little BS Results |  |  |  | Little BS Resources<sup>d</sup> |  |
| --- | --- | --- | --- | --- | --- | --- | --- | --- | --- | --- | --- | --- | --- | --- | --- |
| | Sites (L) | Seqs | Unique Sites per Seq. | *g* | *s* × *r* | Sites (*l*) | Uniq. sites<sup>b</sup> | Uniq. Sites per Seq. | Total Uniq. sites<sup>c</sup> | Avg. *BCL* | $\Delta BCL$ | TPR ($\geq 70\%$) | TPR ($\geq 95\%$) | Time (Hours) | Memory (GB) |
| Butterflies | 5,267,461 | 61 | 61,684 | 0.700 | 4 × 10 | 50,714 | 1.3% | 793 | 5% | 100% | 0.0% | 100% | 100% | 1.37 | 0.38 |
| Plants A | 4,246,454 | 16 | 11,897 | 0.700 | 4 × 5 | 43,614 | 3.3% | 389 | 9% | 100% | 0.0% | 100% | 100% | 0.08 | 0.01 |
| Insects A | 3,011,544 | 174 | 11,758 | 0.700 | 6 × 8 | 34,289 | 1.6% | 188 | 8% | 97% | -1.8% | 98% | 98% | 3.80 | 0.74 |
| Insects B | 2,938,039 | 48 | 29,092 | 0.747 | 10 × 12 | 67,401 | 3.8% | 1,103 | 25% | 91% | -4.5% | 100% | 94% | 3.80 | 0.33 |
| Insects C | 1,719,036 | 193 | 7,346 | 0.748 | 5 × 7 | 46,331 | 3.2% | 236 | 15% | 97% | 1.1% | 96% | 99% | 5.80 | 1.14 |
| Mammals | 1,391,742 | 37 | 20,962 | 0.700 | 7 × 9 | 19,976 | 2.3% | 485 | 13% | 98% | 0.0% | 97% | 100% | 0.32 | 0.11 |
| Spiders A | 137,170 | 27 | 3,071 | 0.800 | 4 × 8 | 12,877 | 13.4% | 411 | 39% | 94% | 0.0% | 96% | 90% | 0.08 | 0.04 |
| Plants B | 135,243 | 30 | 795 | 0.800 | 7 × 9 | 12,732 | 14.2% | 113 | 56% | 99% | -1.0% | 100% | 96% | 0.03 | 0.01 |
| Spiders B | 89,212 | 34 | 1,296 | 0.800 | 9 × 15 | 9,128 | 14.4% | 186 | 66% | 97% | 0.0% | 93% | 90% | 0.06 | 0.03 |
| Birds | 61,794 | 39 | 750 | 0.863 | 6 × 10 | 13,633 | 26.7% | 201 | 80% | 90% | -3.3% | 97% | 93% | 0.11 | 0.04 |

<sup>a</sup>See *Methods* for citations to source studies.
<sup>b</sup>Proportion of unique sites per subsample.
<sup>c</sup>Proportion of unique sites that made into at least one little subsample.
<sup>d</sup>Number of unique sites in all subsamples.
<sup>e</sup>Per little BS replicate dataset.
*g*, *s*, and *r* are the parameters of little BS analysis.
*l*: number of sites in the little subsample ( $l = L^g$ )
*BCL* is the bootstrap confidence limit.
$\Delta BCL$ is the difference between the averages of *BCLs* from little and standard BS.
TPR: True Positive Rate at the given *BCL* cutoff.

### The little BS approach with median‐bagging

We found that little BS with mean‐bagging did not produce 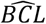 ≥ 95% for 32 species groups, which are false negatives (7.2%) because the standard BS supported all correct species groups at this *BCL* cutoff. These 32 species groups were connected with relatively short branches (< 0.04 substitutions per site; Fig. 2b). Their confidence limits were underestimated by as much as 24% (Fig. 2b). Our investigation into the cause of this underestimation revealed that the distribution of little sample ***bcls*** for these species groups was skewed and that the mean was not the accurate measure of central tendency (Fig. 2c). This prompted us to consider median‐bagging because median is more resilient to outliers. Also, the little BS with median bagging is expected to have the same statistical properties as those established for mean‐bagging^14,19^. However, median bagging has not been previously applied with the bag of little BS in any application in our survey.

The use of median‐bagging eliminated 31 of the false negatives (Fig. 2b), with the remaining species group receiving 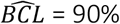 (Fig. 2b). The average 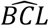 at every branch length cutoff value was greater than 95% for median‐bagging, but not for mean‐bagging (Fig. 2d). We confirmed the improvement offered by median‐bagging for a greater range of *BCL* values by analyzing three gene‐specific sequence alignments (4,000 < L < 10,000; 446 species). Median‐bagging performed much better than mean‐bagging for these short alignments (Fig. 2e) because the distribution of ***bcls*** was skewed and contained many outliers for each dataset (Fig. 2f). Also, false‐negative rates of subsampling approaches become higher when upsampling is not used (Fig. 2g). Therefore, little BS with median‐bagging achieves higher accuracy by overcoming the deficiency of mean‐bagging and traditional divide‐and‐conquer approaches.

### Automatic parameter tuning and the precision of *BCL* estimates

We have developed a simple, automated protocol to determine the three key parameters for little BS analysis (***g, s***, and ***r***). It starts with user‐provided (or default) initial values and increments ***r*** and ***s*** iteratively to generate a stable average 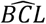 for the whole phylogeny. This step is followed by increasing the size of the little samples by increasing) and re‐optimizing ***r*** and ***s***. This procedure is continued until the average 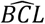 over the whole phylogeny and the number of species groups receiving 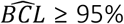 are maximized (see the *Methods* section for details). Its application to 446×134,131 dataset suggested using ***g*** = 0.8, ***s*** = 4, and ***r*** = 6, which confirmed all correct species groups with 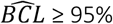 (Average 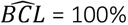). We used this automated system to analyze empirical sequence alignments and discussed its usefulness in the next section (Table 1).

During the automatic determination of little BS parameters, we also estimate the standard error (SE) of 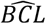 estimates by a procedure in which little samples as well as replicate phylogenies are resampled with replacement (see *Methods* for details). The estimated 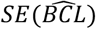 values were inversely proportional to 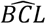 (Fig. 2h). Notably, high precision for 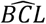 was achieved even when using small ***r*** and ***s*** because 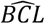 values were very high for most of the species groupings in large datasets.

### Performance of little BS for empirical datasets

We analyzed ten empirical datasets from a variety of species, including eutherian mammals, birds, butterflies, insects, spiders, and plants. The number of taxa in these datasets ranged from 16 to 193, and the number of sites was as many as 5.2 million (Table 1). The accuracy of little BS with median bagging was excellent for all empirical datasets analyzed. The true positive rate (TPR) at 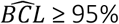 was greater than 95% for six datasets and 90% for the other four (Table 1). TPR at 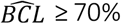 was greater than 95% for nine datasets and 93% for the other one. Phylogeny‐wide average 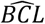 was close to that from standard BS ***BCL***, as the average difference was only 0.1% (Table 1).

The high TPR and 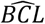 accuracies for larger datasets were achieved by analyzing little subsamples containing only a fraction of all sites (Table 1). Consequently, the memory required to analyze each little BS replicate was tens to hundreds of MBs rather than multiple GBs. The computation time was in minutes or a few hours per little dataset, depending on the number of sequences (Table 1). For example, the little BS analysis of the mammalian dataset (Table 1) required 0.1 GB per replicate, on average, rather than 3.1 GB RAM (∼29‐fold memory savings) and 0.32 CPU hours rather than 9.8 CPU hours per bootstrap replicate (31‐ fold time efficiency). This translated into greater than 95% savings in both memory and time, which enabled us to run many little BS replicates on a standard multicore personal desktop equipped with modest memory (8 GB).

The little BS analysis needed smaller subsamples (smaller ***g***) for empirical datasets with larger numbers of unique site configurations per sequence (C/S; Table 1, and Fig. 2i). Analysis of datasets with fewer than 100,000 unique site configurations required subsamples containing a much larger fraction of site configurations (13%‐27%) than those with millions of site configurations (1.3%‐3.3%). This means that the little subsamples already contained sufficient information for robust phylogenetic inference, as their *C*/S ratios were very large (113 – 1,103) even though an order of magnitude smaller than the full dataset (750 – 61,684; Table 1) in little BS replicate datasets. Interestingly, however, a vast majority of unique site configurations for smaller datasets were included in at least one little subsample, but the opposite was the case for datasets with greater than 100,000 unique sites in our empirical data analysis (Table 1). We found a similar pattern in the little BS analysis of the computer‐simulated 446×134,131 dataset. The use of little samples of size *l* = L^0.8^ and L^0.7^ had C/S of 28.7 and 8.7, respectively, which was more than an order of magnitude smaller than the full data C/S of 300.7. Still, TPR was very similar: 100% and 99.7%, respectively, for *l* = L^0.8^ and L^0.7^. Therefore, little BS analysis with a relatively small C/S value produces results similar to those from standard BS of the full dataset.

### Combining little BS with other optimizations

We also evaluated the performance of little BS when combined with the Ultrafast bootstrap^11^ (UFB). UFB makes standard bootstrapping faster for a large number of sequences. For the mammalian dataset, the Little BS + UFB required only 50 minutes (0.2 GB RAM) on a computer with 5 cores when using ten little samples (*r* = 1,000, default in IQTREE^11^). This was much faster and leaner than using only one of the optimizations: UFB itself required 7.1 GB of RAM and 4.5 hours, whereas little BS alone required 19.8 hours and 0.1 GB of RAM. Therefore, plugging‐in the UFB optimization for generating sample‐wise ***bcl*** ‘s further increases memory and time savings. In the future, we expect little BS to be used along with other efficient heuristics developed to speed up bootstrap calculations^10,11^, and one may use Transfer Bootstrap^2^ when estimating confidence limits.

## CONCLUSIONS

With the rise in large genomic datasets assembled from burgeoning sequence databases, the computational demands of Felsenstein’s traditional bootstrap approach have become a major bottleneck limiting robust and reproducible phylogenetic research. The little bootstraps approach helps remove this bottleneck and enables parallelization even with modest computational resources. Ultimately, computationally efficient approaches will promote greater scientific rigor for all involved in building the tree of life, which requires assessing the robustness of inferences to selecting biologically distinct subsets of data, choice of substitution models and strategies, and application of a myriad of ways of combining multigene datasets.

## METHODS

### Simulated and empirical sequence data assembly

We analyzed multigene alignments assembled from a collection of simulated datasets analyzed in the previous studies^18,21–23^. These datasets were simulated using an evolutionary tree of 446 species (Fig. 2a)^18,24^ A wide range of biologically realistic parameter values derived from empirical data^18^ was used in simulating hundreds of gene alignments, including sequence length (445 – 4,439 bases), G+C content (39 – 82%), transition/transversion rate ratio (1.9 – 6.0), and gene‐wise evolutionary rates (1.35 – 2.60×10^‐6^ per site per billion years)^18,21^. Evolutionary rates were also heterogeneous across lineages, simulated for each gene independently under autocorrelated and uncorrelated rate models^18,21^. Simulated alignments of 100 genes that evolved with the autocorrelated rate model were concatenated to form the 446×134,131 (species x bases) dataset. The 446×536,524 sequence alignment was generated by concatenating sequence alignments generated by concatenating 100 randomly selected gene alignments from each of the four different lineage rate variation models simulated in ref.^18^. Three smaller datasets were analyzed, corresponding to individual simulated genes: 446×4,070, 446×7,002, and 446×9,359 bases.

Ten empirical datasets were analyzed consist of DNA sequence alignments. These datasets consisted of sequences from eutherian mammals^4^, butterflies^25^, plants (A^26^ and B^27^), insects (A^28^, B^29^, and C^30^), spiders (A^31^ and B^32^), and birds^33^ (Table 1). The number of taxa ranged from 16 to 193, and the number of sites ranges from 61,794 to 5,267,461. We used the phylogenetic trees (ML trees) presented in the original studies as the reference trees for empirical datasets. The ground truth for little BS confidence limits were the standard BS confidence limits reported in those published articles.

### Standard and little bootstrap (BS) analyses

We used the IQTREE software^34^ with a general time‐reversible nucleotide substitution model with gamma‐distributed rate variation (GTR+Γ)^35,36^ and default ML search parameters. One hundred replicates of standard bootstrap analyses were conducted to generate ***BCL***s, all of which were very high for large datasets analyzed. For three single‐gene datasets, 1,000 bootstrap replicates were used to generate stable ***BCL***s. The confidence limits obtained using the standard bootstrap analyses were the ground truth in our analyses, as the bag of little bootstraps is being investigated as a computationally efficient alternative. The true tree used in computer simulations was the reference in the analysis of simulated datasets. The bootstrap confidence limits presented in the published phylogenies were used as references for the empirical datasets analyzed. The parameters of little BS analyses for these datasets were selected using the protocol presented below. We also applied the Ultrafast bootstrap (UFB)^11^ on the mammal dataset using (GTR+Γ)^35,36^ model in IQTREE with the default option of 1,000 replicates. For little bootstrap analysis, the UFB with the same options was carried out for each little dataset directly to estimate the required time and memory. These are approximate estimates because IQTREE does not have a provision for upsampling when generating bootstrap replicate datasets. The reported estimates are expected to be very close to the actual time estimates because IQTREE compresses identical site configurations during ML calculations, and upsampling only alters site configurations’ frequencies.

### Automatic selection of little BS parameters

Our procedure automatically determines the size of the subsample (*g*), the number of subsamples (*s*), and the number of replicates (*r*). The procedure starts with *g* = 0.7 if the sequence alignment contains ≥ 100,000 unique site configuration (such that *l* < 50,000), otherwise we set *g* = 0.8. One may set any starting or fixed value of *g*. In step 1, we conduct little BS with *s* = 3 and *r* = 3 to generate initial 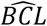 for all the nodes in the given phylogeny (if provided) or from a majority rule bootstrap consensus tree. Using these values, we generate average 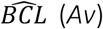 and the fraction of inferred tree partitions with 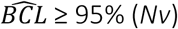 . Through an iterative process, we stabilize and maximize both *Av* and *Nv*, as follows. In step 2, we add one little BS replicate to each subsample (i.e., *r* increases by 1) and then compute *Av*. We repeat steps 2 and 3 by increasing *r* until the difference in successive *Av* values is less than 0.1% (or a user‐specified threshold, δ_r_). In step 4, we increase *s* by one and generate *r* additional replicate datasets and phylogenies, and compute *Av* and *Nv*. If the difference between *Av* for the current (*s*) and the previous (*s* ‐1) set of subsamples is greater than 1% (or user‐specified δ_s_), then we repeat step 4. In step 5, we check and see if *Nv* is less than 100% or the *SE* of estimated 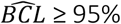 is too high (>5%). If so, we increase the little subsample size by and restart the analysis from step 2. In step 6, we go to step 4 if the user‐specified precision (SE) has not been achieved.

### Estimating standard errors of 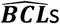

Given ***r*** bootstrap replicate‐phylogenies for ***s*** samples, we employ a bootstrap procedure to generate SE of 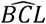. We use already computed phylogenies of *r* × *s* little BS replicates and derive 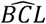 for all the nodes from collections of phylogenies by resampling ***s*** samples with replacement and ***r*** replicates with replacement every time a subsample is selected. This process is repeated 100 times, and the standard deviation of each tree partition’s 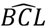 is generated to estimate its *SE*. This process is extremely fast because precomputed phylogenies are used.

### Phylogenomic subsampling approaches without upsampling

We also generated 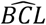 values by a little BS procedure in which upsampling was replaced by the standard BS resampling such that the replicate datasets contained only *l* sites rather than ***L*** sites. We refer to this as the Phylogenomic Subsampling with Resampling (PSR) approach, in which one may use either mean‐ or median‐bagging. We also generated 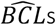 without any resampling or upsampling (i.e., ***r*** = 0) such that the ML phylogenies were inferred from ***s*** subsample datasets containing ***l*** sites each. We call it the Phylogenomic Subsampling (PS) approach. We compared the true positive rates 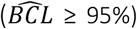 of little BS, PSR, and PS approaches for the computer‐ simulated 446×134,131 dataset (***g*** = 0.7) For all analyses, 100 replicate phylogenies were generated by using ***s*** = 10 and ***r*** =10 for little BS and PSR, and ***s*** = 100 for the PS approach.

### Analysis pipeline for little BS

We developed an R^37^ pipeline to conduct little bootstraps analysis by using IQTREE. In this case, we used the Biostrings^38^ package to generate little datasets of the specified lengths (***l***) and then bootstrap replicate datasets in which L sites were resampled with replacement from ***l*** sites. The resulting datasets were used to obtain ML phylogenies that were summarized by using the function *plotBS* from the phangorn^39^ library that produced the ***bcl*** for each of the phylogenetic groups in the standard bootstrap phylogeny. Mean and median‐bagging estimates were obtained from sample‐wise ***bcl***s from little samples using a customized function in *R*. We used ten samples and ten bootstrap replicates for little bootstraps analysis for concatenated gene datasets, and 50 little samples and 20 bootstrap replicates for single‐gene datasets. We applied the automated protocol using a customized R function. We also developed a customized R function for estimating SEs of 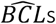.

## Data availability

Datasets analyzed are available from https://doi.org/10.6084/m9.figshare.14130494.

## Code availability

R codes are available from https://github.com/ssharma2712/Little-Bootstraps. A capsule containing source codes and datasets for our analyses is available on the CodeOcean service (https://doi.org/10.24433/CO.6432188.v1). Users can replicate the little bootstraps sampling and bagging steps in this capsule.

## Acknowledgments

We thank Sara Vahdatshoar and Julia Davis for their help with computational analysis. We thank Drs. Jack Craig, Qiqing Tao, Marcos Caraballo‐Ortiz, Antonia Chroni, Cristian Palacios, Sergei L. K. Pond, and S. Blair Hedges for providing critical comments on the manuscript.

## Author Contributions

S.K. conceived all the methods, designed analyses, developed visualizations, and wrote the manuscript. S.S. refined methods, designed and conducted analyses, refined visualizations, and contributed to writing the manuscript.

